# Matching provides efficient decisions

**DOI:** 10.1101/2024.02.15.580481

**Authors:** Jan Kubanek

**Affiliations:** University of Utah, Salt Lake City, Utah, United States

## Abstract

How humans and animals distribute their behavior across choice options has been of key interest to economics, psychology, ecology, and related fields. Neoclassical and behavioral economics have provided prescriptions for how decision-makers can maximize their reward or utility, but these formalisms are used by decision-makers rarely. Instead, individuals allocate their behavior in proportion to the worth of their options, a phenomenon captured by the generalized matching law. Why biological decision-makers adopt this strategy has been unclear. To provide insight into this issue, this article evaluates the performance of matching across a broad spectrum of decision situations, using simulations. Matching is found to attain a high or near-optimal gain, and the strategy achieves this level of performance following a single evaluation of the decision options. Thus, matching provides highly efficient decisions across a wide range of choice environments. This result offers a quantitative explanation for the broad adoption of matching by biological decision-makers.

## Introduction

How humans and animals allocate behavior, time, and effort to their options has been a fundamental question in science and human society. Neoclassical economics provides a normative apparatus that shows how humans and animals can allocate their time, effort, or monetary resources [1, 2]. A hallmark of this normative set of theories is that decision-makers maximize a criterion—individuals allocate their resources to maximize their total utility or payoff [3–5]. Despite its normative appeal, this framework is adopted by decision-makers infrequently [6, 7].

Biological sciences have reported a much more common strategy in which decision-makers allocate their behavior in proportion to the value of their options [8–13]. This behavior has been observed so commonly that it has been formalized as the matching law [8–10, 12, 14, 15]. When generalized—allowing for the representation of value to be nonlinear [16–23]—the matching law explains much of choice behavior, including undermatching, overmatching, and their specific forms [15, 19, 24–26].

Why decision-makers adopt matching so frequently has been a conundrum to decision sciences, in particular because matching is generally suboptimal [2, 15, 27–29]. Decades-long debates have centered on whether matching might represent a universal heuristic to optimal choice behavior [13, 15, 30–32]. This issue has been difficult to resolve because it is not known how matching performs at a general level and what its specific strength is.

Previous studies assessed the performance of matching in specific tasks, particularly focusing on variable-interval and variable-ratio schedules [8–10, 12, 14, 15]. An analytical study attempted to evaluate the performance of matching at a general level [13], but the analytical approach can only identify the necessary—and not the sufficient—contingencies between utility and effort for which matching is optimal.

To address this issue, this article uses a computational approach to evaluate the performance of matching at a general level. Using this approach, matching is also assessed along decision time, i.e., the necessary number of evaluations of decision options. This methodology uncovers a striking finding: Matching harvests on average around 90% of the total available reward following a single evaluation of the worth of decision options.

## Results

### Assessment of matching efficiency

This article assesses the performance of matching over a broad spectrum of decision situations. The evaluations include 2, 3, and up to 10 decision options, each with distinct contingency between relative utility and relative effort (Materials and Methods). These contingencies include reinforcement schedules that have been used to assess the performance of matching in previous studies [8–10, 12–15], and generalize the spectrum to many other contingencies and number of options, for a total of 9000 decision settings (Materials and Methods). In each case, the performance of matching is quantified relatively to maximization, which provides the highest possible gain. The performance of matching and maximization is further assessed over the number of evaluations of the decision-makers’ options.

Figure 1 shows the principal finding. Matching decision-makers (green) obtained an average 90.2% of the total possible gain across all choice situations. Notably, they collected this level of gain following a single evaluation of the worth of each option. In contrast, maximizing decision-makers required an average of at least 16 evaluations to reach equivalent performance (Figure 1, gray).

**Fig 1.**
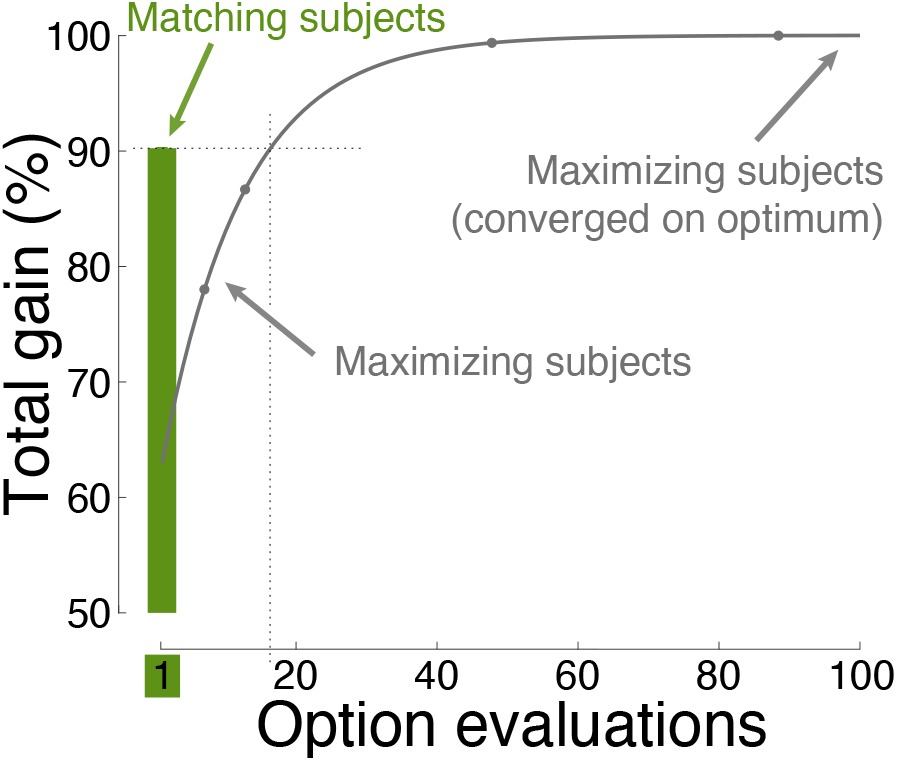
Convergence on optimal choice behavior. The performance of matching and maximization as a function of the number of evaluations of the choice options. The assessment was performed over 9000 choice situations, involving up to 10 options, each with distinct contingencies between reward and effort (Materials and Methods). The maximum gain (the value of 100%) was established by allowing maximization to converge. Matching (green) provides high gain following just one sample (one evaluation; see abscissa) of the worth of the choice options. A specific example of how matching achieves high performance following a single evaluation of the choice options is shown in Figure 3B. Here and henceforth, plots show mean*±* s.e.m. values. Here and elsewhere, the error bars are often smaller than the symbols indicating the mean.

To exemplify this finding, consider a foraging animal that must decide how to allocate her presence across several patches of food. Matching allows her to visit each patch just once to evaluate its worth, and allocate her presence across the patches in proportion to their relative worth. If she maximized reward, she would have to visit each patch at least 16 times to converge on a behavioral allocation of quality comparable to matching.

Thus, matching allows individuals to make effective decisions in a single step, i.e., matching provides highly efficient decisions.

### Efficiency of generalized matching

Matching can explain much of choice behavior when generalized [16–20]. The following analysis investigates the efficiency of generalized matching, i.e., performance following a single evaluation. In generalized matching, the relative utilities of choice options are parameterized using an exponent *β* (Materials and Methods). This exponent implements tendencies for undermatching (*β <* 1) or overmatching (*β >* 1). Specifically, decision-makers who undermatch the relative utility proportions are relatively indifferent to reward. Decision-makers who are drawn to rewarding options overmatch the relative utility proportions. In the extreme, highly over-matching, greedy decision-makers put all their effort into the seemingly richest option (*β »* 1).

Figure 2A shows the performance of generalized matching as a function of *β* across the space of choice situations. The figure reveals that *β* = 1.0, which characterizes strict matching [33–35], lies near the optimum, with *β* = 1.0 and *β* = 1.2 delivering a 90.2% and 90.5% gain, respectively. Overmatching and undermatching are similarly effective in their typical, mild forms [25, 26], but incur a performance loss when extreme. In the extremes, the greedy strategy, compared with indifference, provides a relatively high gain over an appreciable range of *β* values (Figure 2A). The following analyses therefore compare the performance of matching also with this degenerate form.

**Fig 2.**
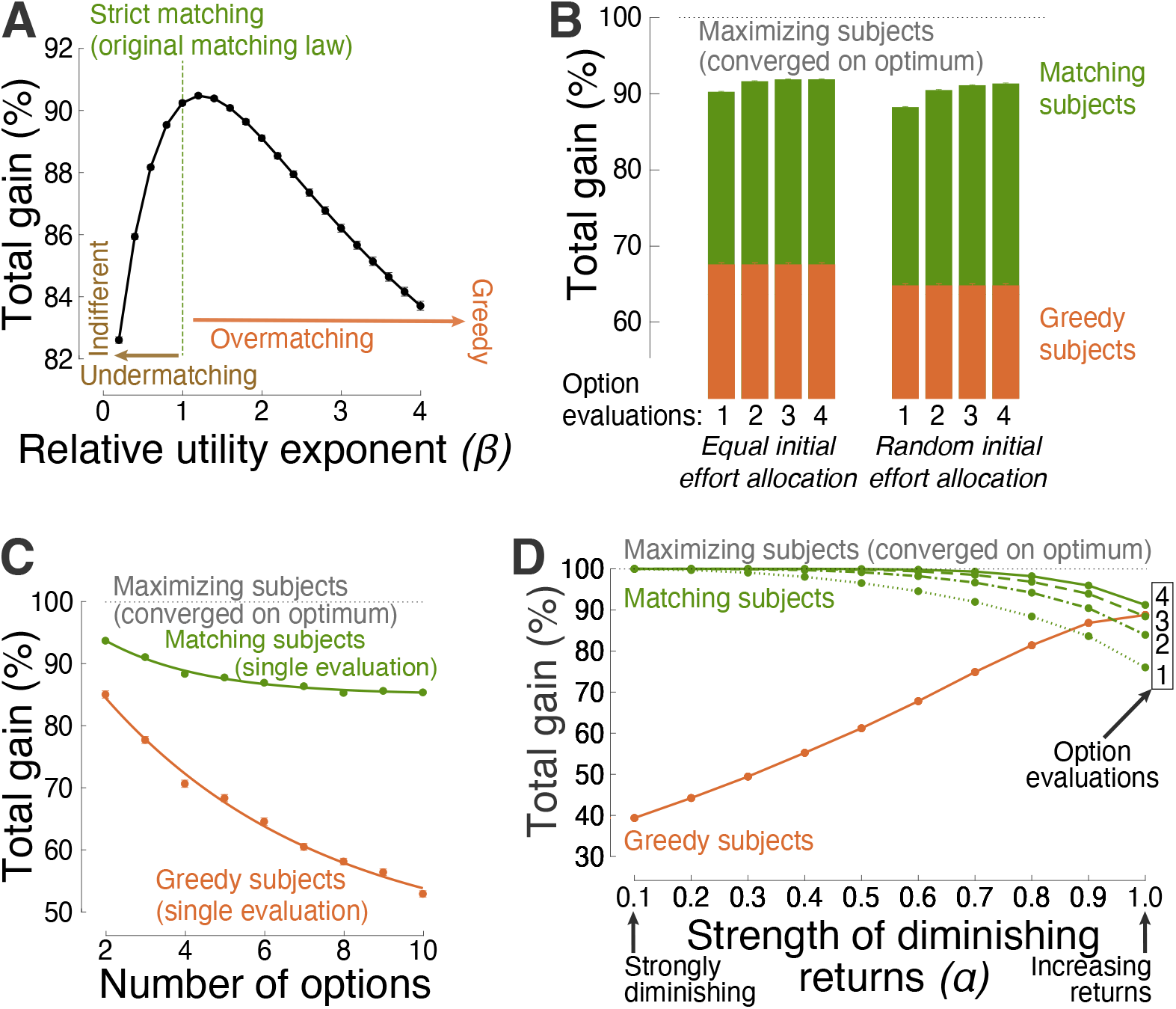
Efficiency of matching across choice situations. (**A**) Gain as a function of the level of the exponent *β* in the representation of relative utilities (Materials and Methods). Strict matching of the original matching law [33] uses *β* = 1, i.e, relative effort is allocated in proportion to relative utilities that are not exponentiated. In this figure, the same 9000 decision situations as in Figure 1 were used. (**B**) Performance comparison associated with initial effort allocation to assess the initial worth of decision options. The left (right) bars correspond to equal (random) allocation of initial effort. This more general, random initial distribution is used henceforth. Matching decision-makers may sample the worth of their options repeatedly 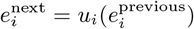; see Figure 3B for a specific example). The individual bars show performance as a function of the number of the repeated option evaluations (same abscissa as in Figure 1). Same data as in Figure 1, separately for a specific number of decision options. The greedy strategy is superimposed for comparison. The curves represent exponential fits. Performance as a function of the level of diminishing marginal utility. To evaluate the full spectrum from strongly diminishing to not at all diminishing (linearly increasing) returns, this analysis specifically used the power contingency between utility and effort (see Materials and Methods). The abscissa shows performance as a function of 10 evenly spaced values of the power *α*. Like in **B**, the decision strategy 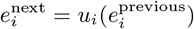 can be applied repeatedly. Performance curves for up to 4 option evaluations are shown in distinct line styles.

### Efficiency across choice situations

In changing environments, decision-makers may apply maximization or matching repeatedly [36–40]. For matching, this repeated application, i.e., 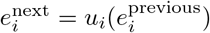, is an instance of melioration [41]. Melioration is shown to increase the performance of matching further. Figure 2B, left green bars, shows that a second evaluation and effort allocation increases the average performance from 90.2% to 92.0%. The performance saturates in about the third and fourth evaluation, reaching 92.3% of the possible maximum. The performance of the greedy strategy does not, by definition, improve with additional evaluations (Figure 2B, left orange bars; see also Figure 3B for a specific example).

**Fig 3.**
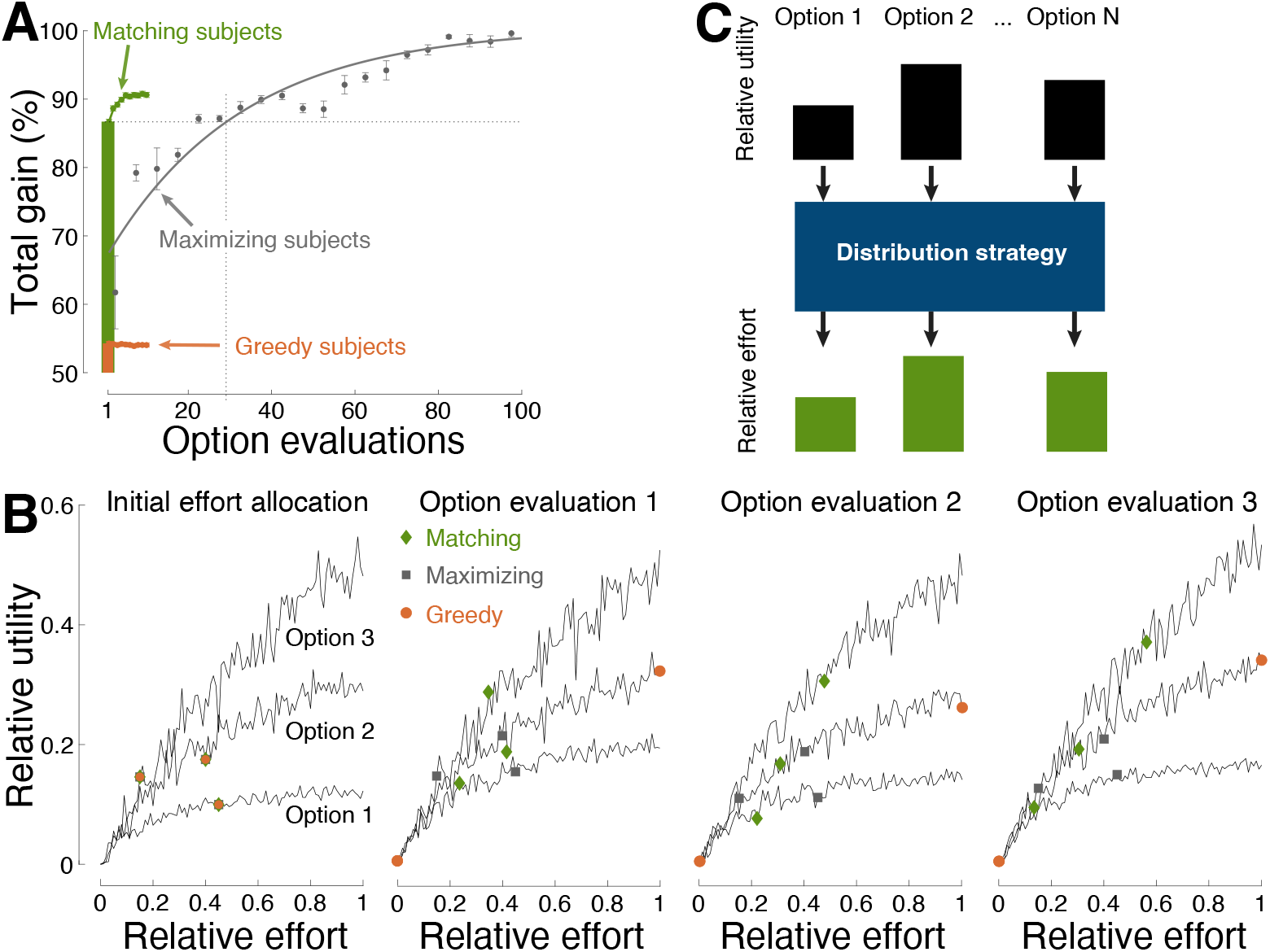
Efficiency in stochastic environments. (**A**) Same format as in Figure 1 for choice situations in which each option provided a distinct rate of reward, according to variable interval schedules (Materials and Methods). Performance is shown over 250 (10) consecutive evaluations of the decision options for maximizing (matching) decision-makers. The strategies initiated with a random allocation of effort across the choice options. (**B**) Example of effort allocation in a three-option situation across individual evaluations. The evaluation steps associated with the individual strategies are shown as colored symbols (see inset). All strategies started with the same random initial allocation of relative effort. (**C**) Summary. This study focused on the distribution strategies (blue box) that individuals use to allocate relative effort to the relative utilities associated with their options. A proportional (Figure 2A) allocation of relative effort to relative utilities (i.e., matching) is found to provide rapid convergence on optimal decisions.

Thus far, decision-makers were assumed to allocate effort equally when sampling the initial worth of their options (Materials and Methods). This case also incorporates situations in which individuals can estimate the worth of their options by direct means (e.g., price), in which no initial sampling is required. However, in many cases in nature, humans and animals must sample their options to learn about their initial worth. Equal sampling may be impractical or impossible. The following analysis reveals that the efficiency of matching is maintained regardless of the type of initial distribution of effort. Specifically, Figure 2B, right green bars, shows that decision-makers who distribute their initial effort entirely randomly (e.g., Figure 3B) can still apply the same strategy with comparable level of efficiency. In addition, the performance of matching increases even more steeply by re-adjusting effort in response to additional evaluations 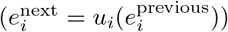. On the fourth evaluation, the performance reaches 91.6%. A two-way ANOVA with factors evaluation number (*I*) and initial effort distribution (*D*) detected a significant effect of *I* (*F* (3, 71992) = 672.42, *p* = 0), a significant effect of *D* (*F* (1, 71992) = 715.82, *p* = 6.4 *×* 10^*−*157^), and a significant interaction between the factors (*F* (3, 71992) = 51.38, *p* = 3.6 *×* 10^*−*33^). The greedy strategy does not depend on the number of evaluations (Figure 2B, orange) but is sensitive to the initial allocation of effort (*F* (1, 71992) = 300.10, *p* = 4.3 *×* 10^*−*67^).

Matching maintains its efficiency also in regard to the number of decision options. Figure 2C shows that matching is efficient even when faced with a relatively high number of options. Following a single evaluation, matching attains an average 93.7% gain for all choice situations that involved 2 options. The strategy has an exponential decay of 2.6% per option, and achieves a 85.3% gain for 10-option decisions. In comparison, greedy subjects fall short when facing an appreciable number of options. This is because putting all effort into the seemingly most rewarding option leads to an opportunity loss (e.g., Figure 3B). The greedy strategy shows an exponential decay of 3.9% per option, and attains a mere 52.9% average gain in 10-option situations.

Matching is found to be particularly efficient in situations with diminishing returns, but provides high performance also in situations in which the law of diminishing marginal utility does not hold. To quantify how diminishing-increasing returns affect matching performance, the strength of diminishing returns is parameterized using the power function, *U* (*E*) = *U*_*m*_*E*^*α*^, where *α* is the tested parameter. For *α* = 0.1, the gain *U* diminishes strongly with invested effort *E*. On the other hand, for *α* = 1.0, gain increases linearly with effort and never saturates. Figure 2D shows the dependence of the total decision gain on this parameter. Matching is highly efficient—approaching perfect gain following a single evaluation of the decision options—when the choice options obey the law of diminishing marginal utility. This reproduces a previous, analytical study [13]. The degenerate, greedy strategy is particularly inadequate in this case: A greedy individual puts all effort or resources into an option whose payoff eventually saturates, thus experiencing an opportunity loss. As expected, the performance of matching/greedy behavior is lower/higher in choice situations in which the law of diminishing marginal utility does not hold. Nonetheless, matching can outperform greedy behavior also in these cases, when applied repeatedly 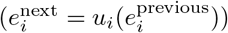. In particular, following the fourth evaluation, matching attains a 91.2% gain, compared to a 88.8% gain of the greedy approach. The same principal observation is made for functions other than the power function (Figure 4B).

**Fig 4.**
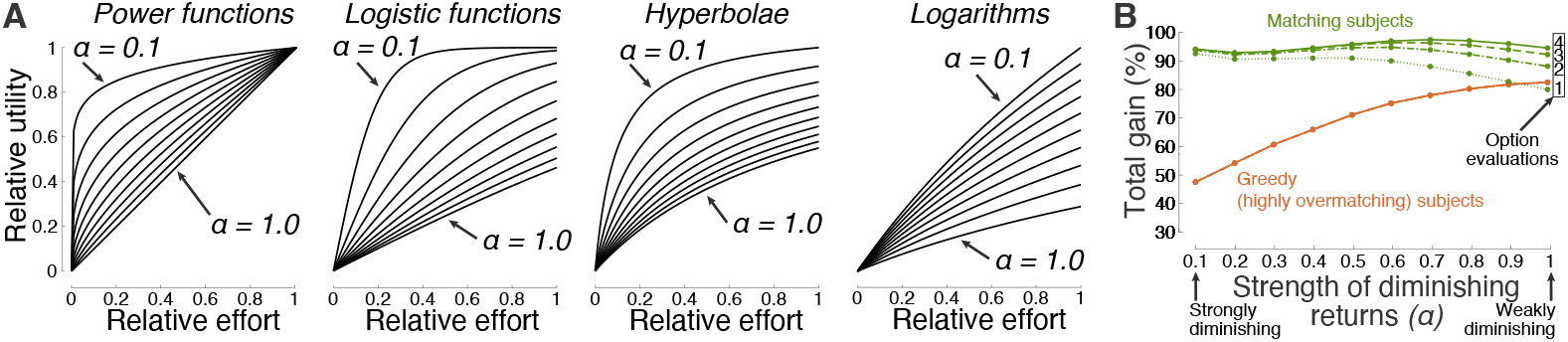
Choice environments and the effect of diminishing marginal utility. Choice environments tested. The study evaluated the performance of matching over a large space of utility-effort contingencies. Power, logistic, hyperbolic, and logarithmic functions in the forms 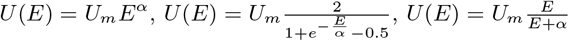, and 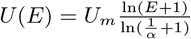, respectively, were used. These contingencies include reinforcement schedules that have been used to assess the performance of matching in previous studies [8–10, 12–15], and generalize the spectrum to many other contingencies and number of options, for a total of 9000 decision settings (Materials and Methods). In these functions, the variable *α* quantifies the level of diminishing marginal utility, and ranged from 0.1 (strongly diminishing) to 1.0 (least-diminishing; linear for the power function), in steps of 0.1. The individual plots are scaled by these 10 values of *α* in the indicated order. In addition, the variable *U*_*m*_ in these functions scales the magnitude of each contingency, ranging from 0 to 1. (**B**) Average performance of matching and its greedy counterpart as a function of the level of diminishing marginal utility. Same format as in Figure 2D, but now for any combination of the functions shown in **A**. Specifically, each option (2 up to 10; 1000 evaluations each) was randomly assigned any of the four functions with the diminishing marginal utility level *α* indicated on the abscissa. Strong diminishing returns provide an almost immediate convergence onto the optimum. Iterative application of matching greatly improves performance in cases with weakly diminishing or increasing returns (see number of option evaluations in the inset).

### Efficiency in stochastic environments

Matching provides a high-level strategy to effort allocation. Therefore, the results presented thus far are general to any task or choice situation that provides a certain kind of reward as a function of a certain kind of effort. This includes deterministic and stochastic choice situations. As a specific example of performance in stochastic choice environments, matching was deployed in environments in which rewards were provided according to the frequently investigated variable interval schedules [8, 13, 14, 33–35, 38, 42–47]. Matching and maximizing decision-makers integrated the individual binary rewards over a defined period of time using the commonly used reinforcement learning [38, 39, 48], and allocated effort according to the estimated utility of each option (Materials and Methods). The rate of reward occurrence for each option was again randomized across 1000 distinct choice situations, each involving 2, 3, and up to 10 options.

Matching was found to be rapid and efficient also in this case (Figure 3A). Following a single evaluation, matching harvested an average of 86.7% of the total possible gain. In comparison, maximization required an average of 29 option evaluations to reach equivalent performance. The greedy strategy produced a mere 54.1% gain.

Figure 3B exemplifies how matching achieves its high efficiency. Following an initial assessment of the relative worth of the available options (left panel), matching (green diamonds) allocates effort in proportion to the relative worth, i.e., 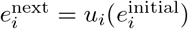. The second panel shows that this step readily results in harvesting the highest reward from the richest option (i.e., Option 3), as desired, while also gaining substantial reward from the relatively less valuable options. Repeated applications of matching (third and fourth panels) lead to additional gain. Maximization (gray squares) is found to require a large number of option evaluations to reach the performance level of matching (Figure 3A). The degenerate, greedy strategy (orange circles) settles with the option that appear richest at that moment, and thus misses the globally richest option.

## Discussion

This study finds that matching provides effective behavioral allocations across a broad spectrum of decision situations, yielding high or near-optimal gain. Furthermore, the strategy is found to possess a notable strength—high efficiency. Matching approaches near-optimal gain following a single evaluation of the worth of the decision options (Figure 1, Figure 2, Figure 3). This finding holds across distinct choice situations (Figure 1), generalized forms of matching (Figure 2A), initial allocation of effort (Figure 2B), number of options (Figure 2C), the level of diminishing returns (Figure 2D), and stochastic environments (Figure 3).

The article finds that matching harvests on average around 90% of the maximum possible gain following a single evaluation of the worth of the decision options. Moreover, in economic settings, in which the worth of the options may be known or can be estimated (e.g., price), matching allows individuals to allocate effort (e.g., money) readily, without any prior sampling. Thus, matching is efficient, combining high performance with rapid evaluation.

The time to make a choice is a factor crucial to survival and rewarding behavior [49–53]. Indeed, most decisions involve a speed-performance or speed-accuracy tradeoff [54, 55]. This natural constraint limits the number of times individuals can assess and evaluate their options. Reward maximization, which provides the highest gain possible, is found to be taxing in this regard. For instance, Figure 1 and Figure 3 show that reward maximization, even when algorithmically efficient (Materials and Methods), would require a decision-maker to visit each option or food patch on average at least 16 times to make a decision of quality comparable to matching. In comparison, matching provides a single-step convergence on the optimum (Figure 1, Figure 2, Figure 3). Individuals using this efficient decision strategy may possess an evolutionary advantage. Indeed, matching has been considered a candidate for an evolutionarily stable strategy [36, 37]. Referred to as the relative payoff sum in behavioral ecology, a population of individuals using this strategy is believed to be uninvadable by a mutant with a different strategy [36, 56]. A proof was attempted [36] but remains elusive [57]. The present study shows that this strategy is efficient, providing optimal or near-optimal gain after just one sampling of the worth of the choice options. Given that foraging efficiency constitutes an important optimization criterion in ecology [49, 50, 58], the result of this study supports the proposition that this strategy might be evolutionarily stable.

Matching is applicable both at the global level of the matching law and at the local level of decision strategies [38–40, 59]. For instance, matching was found to be used as a time-by-time decision strategy by non-human primates [38, 60]. The subjects allocated their relative effort—measured by the relative number of choices of an option—to the relative utility of each option. Moreover, the relative utility associated with the matching equation, termed fractional income, captured firing rates of neurons in the recorded brain region, the lateral intraparietal area. At the molecular level, matching is known to result from melioration [41], a strategy in which decision-makers gradually shift effort to the more valuable options. The repeated, iterative application of the matching rule shown here, i.e., 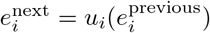 (Figure 2B,D; Figure 3A,B), is an instance of melioration.

Humans and animals often exhibit under- or over-matching tendencies [25, 26]. The analyses performed here find that these slight deviations from the original, strict matching law [33] incur only a minor loss in performance (Figure 2A). Therefore, the principal discovery—the efficiency of matching as a strategy—also applies to the generalized form of matching. Generalized matching allows for the representation of value to be nonlinear, which is supported by several fields [21–23]. In this context, the article also shows that extreme forms of undermatching (i.e., indifference) and overmatching (i.e., greedy behavior) result in relatively low gain and thus can be considered maladaptive.

Decision-makers are known to weigh benefits against costs, or rewards against effort. Yet, how effort should be incorporated in current models of decision-making in economics [61], ecology [62], and psychology [63, 64] has been unclear. This article shows that individuals can make efficient cost-benefit decisions by adopting matching, i.e., evaluating the relative reward *u* per relative effort *e* of each option (in matching, 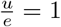 for all options). This value formulation increases with the benefit of an option (*u*), decreases with the cost of an option (*e*), and the relativistic expression does not require absolute units and can thus be represented by neurons.

This study focused on the principles that individuals use to allocate their effort to their choice options (blue box in Figure 3C). The input to these choice rules—how subjects represent the worth of their options—is a separate question that has been addressed in neuroscience [38, 39, 48, 65]. Notably, in matching, the relativistic nature of *u*_*i*_ and *e*_*i*_ enables these variables to be subjective. So long as the underlying representations are comparable by the brain (e.g., through firing rates), the relativistic *e*_*i*_ = *u*_*i*_ strategy provides a rapid and efficient allocation of the decision-maker’s resources.

In matching, the relativistic representation of utility involves divisive normalization. Divisive normalization is a common operation performed by neural circuits [66]. The specific form of this operation may be crucial for explaining attraction, similarity, and compromise effects observed in multi-alternative, multi-attribute decision environments [67, 68]. For instance, it has been found that a transformation of utilities by specific monotonic functions prior to divisive normalization can explain these behavioral effects parsimoniously [68]. Monotonic transformations and divisive normalization are performed by several kinds of feedforward and feedback neural circuits [66, 69].

In summary, this article evaluated the performance of matching across a wide range of choice environments, and uncovered a notable strength of this strategy across the environments—high efficiency. These quantitative findings may explain why biological decision-makers adopt matching so commonly.

## Materials and Methods

### Choice behavior

In general, decision-makers allocate their total effort, 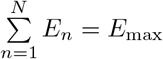, to their *N* options based on the options’ relative utilities or rewards. The effort *E* can represent time, money, a physical or mental expenditure; generally, any resource that an individual allocates to her options in a given choice situation.

#### Maximization

Maximizing decision-makers allocate effort across their choice options to maximize the total reward or utility 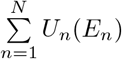 across their options.

#### Matching

In strict matching, decision-makers allocate their relative effort associated with each option *i*, 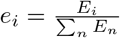 in proportion to the associated relative utility or reward, 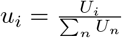, such that *e*_*i*_ = *u*_*i*_ (Figure 3C). Generalized matching also follows *e*_*i*_ = *u*_*i*_, but the relative utility of each option is parameterized as 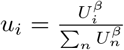, where the exponent *β >* 0. This parametrization captures subjective representation of utility [21–23].

### Evaluations

#### Choice situations

The evaluations sampled a broad spectrum of utility-effort contingencies, i.e., *U* (*E*) functions. The functions were of distinct kinds, used distinct magnitudes, and distinct level of diminishing marginal utility (Figure 4A). Specifically, power, logistic, hyperbolic, and logarithmic functions in the forms *U* (*E*)= *U*_*m*_*E*^*α*^,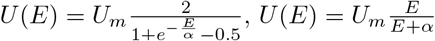, and 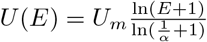, respectively, were evaluated. The logarithmic functions used *E* + 1 in their argument to avoid negative utility, and ln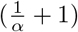 in the denominator to implement logarithm base values 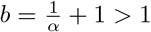. These reward-effort contingencies are encountered frequently in natural and societal settings [13]. The principal findings of this article are relatively insensitive to the particular functions evaluated.

#### Choice options

The study simulated a large number of choice situations to provide broadly applicable results. The situations featured *N* = 2, 3, … 10 options. For each option within the set of *N*, the specific *U* (*E*) function (power, logistic, hyperbolic, or logarithmic), its magnitude *U*_*m*_, and the saturation parameter *α* were all selected randomly. The function kind was selected from the set of the four. *U*_*m*_ was selected randomly from a uniform distribution over the range from 0 to 1. The diminishing marginal utility level *α* was selected randomly from the set *{*0.1, 0.2, 1.0*}*. The randomization was performed 1000 times for each *N*, yielding a total of 9000 choice situations. This number of evaluations provided tight standard errors of the mean effects (Figure 1, Figure 2).

#### Stochastic choice situations

The stochastic environments used the same number of repetitions and the same number of options. Rewards were provided according to the commonly used variable interval schedules [8, 13, 14, 33–35, 38, 42–47]. In a variable interval schedule, a binary reward is assigned to an option randomly from the Poisson process at a specific reward rate [13]. The reward rate was again assigned randomly to each option, within the range from 0 to 1 per unit of time. A reward assigned to an option is available until the decision-maker harvests it. Harvesting events (i.e., the decision-maker’s behavior) are also binary and assigned to an option randomly from the Poisson process, now at the rate of the specific rate of effort (range from 0 to 1 per unit of time, subject to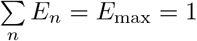). To estimate the reward rate from the noisy binary information, decision-makers must integrate the history of past rewards. A common approach for doing so is reinforcement learning, in which the integrated binary rewards are discounted exponentially [38, 39, 48], i.e., using a function *Ae*^*−γt*^. The learning rate *γ* is related to the time period *T* of integration. This study used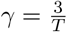, for *T* = 400. The factor *A* was set such that the integral of the exponential over *T* was equal to one, i.e.,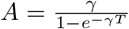. Each option evaluation operated on subsequent time period *T*; i.e., each option evaluation encountered a distinct distribution of rewards and behaviors, generated using the respective rates.

#### Maximization

In each of the choice situations, the maximizing subjects aimed to maximize their gain within a constrained distribution of effort devoted to each option. Specifically, the decision-makers maximized 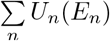 subject to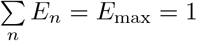. The solution was provided using constrained nonlinear multivariable maximization (function fmincon in matlab, using the default interior-point algorithm; the specific form of the algorithm had only minimal effect on the results). The algorithm converged onto a solution in a finite number of options evaluations in each decision case (see Figure 1 and Figure 3A for the number of evaluations necessary to reach specific performance).

#### Matching

Matching decision-makers also distributed their effort *E*_*i*_ across the choice options subject to finite effort,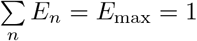. Their total gain,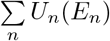,is reported as a percentage of the maximum possible gain harvested by maximization. It was found that matching individuals can use a single or just a small number of evaluations of the decision options. Specifically, they first sampled the worth 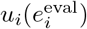 of their options *i* by allocating initial effort 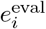 to each option. Subsequently, they applied the matching strategy i.e.,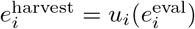. Thus, the decision-makers simply allocated their effort in proportion to an initially established worth of their options. In generalized matching, the proportionality still holds; nonlinearities are modeled using the exponent *β* in the representation of relative utilities (see the Matching paragraph above). In Figure 2B, Figure 2D, and Figure 3, this strategy was applied iteratively using melioration, i.e.,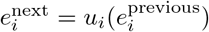. Moreover, the analysis of Figure 2B also tested whether the strategy is sensitive to the kind of the initial effort distribution—equal 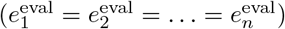 or entirely random.

## Acknowledgements

I thank Drs. Alex Kacelnik, Alasdair Houston, Leo Sugrue, and Julian Brown for comments on the manuscript. This work was supported by the NIH grant RF1NS128569 and NSF grant 2325125. The author has no conflict of interest.

## Code availability

The simulation code is available and documented at neuralgate.org/download/EDM.

## Notes

### Competing Interest Statement

The authors have declared no competing interest.

